# Rapid profiling of drug-resistant bacteria using DNA-binding dyes and a nanopore-based DNA sequencer

**DOI:** 10.1101/2020.10.09.332585

**Authors:** Ayumu Ohno, Kazuo Umezawa, Satomi Asai, Kirill Kryukov, So Nakagawa, Hayato Miyachi, Tadashi Imanishi

## Abstract

Spread of drug-resistant bacteria is a serious problem worldwide. We thus designed a new sequence-based protocol that can quickly identify bacterial compositions of clinical samples and their drug-resistance profiles simultaneously. Here we utilized propidium monoazide (PMA) that prohibits DNA amplifications from dead bacteria, and subjected the original and antibiotics-treated samples to 16S rRNA metagenome sequencing. We tested our protocol on bacterial mixtures, and observed that sequencing reads derived from drug-resistant bacteria were significantly increased compared with those from drug-sensitive bacteria when samples were treated by antibiotics. Our protocol is scalable and will be useful for quickly profiling drug-resistant bacteria.

## Introduction

Since the discovery of penicillin by Alexander Fleming in 1928, numerous infectious diseases have been cured by various antibiotics. However, many drug-resistant bacteria, such as penicillin-resistant *Streptococcus pneumoniae* and methicillin-resistant *Staphylococcus aureus*, appeared and prevailed worldwide (1). Moreover, multidrug-resistant bacteria have been detected against several drugs, such as *Pseudomonas aeruginosa* (MDRP) (2), *Acinetobacter baumanii* and super-multidrug-resistant tubercle bacillus (extensively drug-resistant tuberculosis) (3-5), rendering difficulty of antimicrobial treatments. Thus, bacterial detection and appropriate use of antibiotics are very important to prevent the emergence and spread of drug-resistant bacteria.

Conventionally, for the diagnosis of bacterial infectious diseases, bacterial detection and antimicrobial susceptibility testing (AST) are performed using culture-based methods following the guidelines of the Clinical and Laboratory Standards Institute (CLSI) (6, 7). However, based on the CLSI, it takes at least a few days for bacterial identification by AST, and has a limitation due to a low culture-positive rate. Therefore, treatment of bacterial infections at initial diagnosis inevitably depends on empirical approach. Recently, molecular methods using nucleic acids for the detection of pathogens have been developed; however, these methods often result in detection of non-target of interests due to contamination of indigenous or inviable bacteria, hence making meaningful interpretation of the results difficult.

Therefore, a rapid method for the identification of pathogens and their antimicrobial spectra is necessary in addition to antimicrobial stewardship, in order to treat individual infected patients properly and to prevent the spread of drug-resistant bacteria, as a control measure of nosocomial infection. To achieve this purpose, we incorporated two kinds of technologies. One is the propidium monoazide (PMA) that is effective for live/dead cell discrimination (8, 9). PMA is a DNA-intercalating dye with an azide group that makes a covalent bond with a DNA when exposed to bright visible light (absorbance at 465–475 nm) (10). DNAs bound with PMA cannot be amplified by PCR. Since DNA is usually in the cells, PMA cannot interact with them; however, for the dead cells, DNAs are exposed to the outside and can be bound with PMA by irradiation of bright visible light. Therefore, the fraction of DNAs from live/dead bacterial cells differs greatly through PCR amplification, which can be detected by DNA sequencing. The other technology is the rapid 16S rRNA metagenome sequencing. We previously developed a portable system for rapid 16S rRNA metagenome analysis using the nanopore DNA sequencer MinION and laptop computers (11-13). By combining these two technologies, we aimed at establishing a rapid and accurate diagnostic technology that can detect drug-resistant bacteria.

## Results

### Antimicrobial sensitivity testing (AST)

It has been reported that PAO1 exhibits drug resistance to ampicillin and MDRP exhibits drug resistance to ampicillin and gentamicin (14), but we confirmed the reactivity of antibiotics in actual bacteria. The bacterial suspension (10^7^ CFU/mL) was dispensed onto the plate, and then ampicillin or gentamicin was added. After culturing, the quantity of *E. coli* was decreased with ampicillin and gentamicin treatment, and the quantity of PAO1 was decreased with ampicillin treatment. In contrast, the quantity of MDRP did not change by antibiotic treatments (Figure 1A).

**Figure 1.**
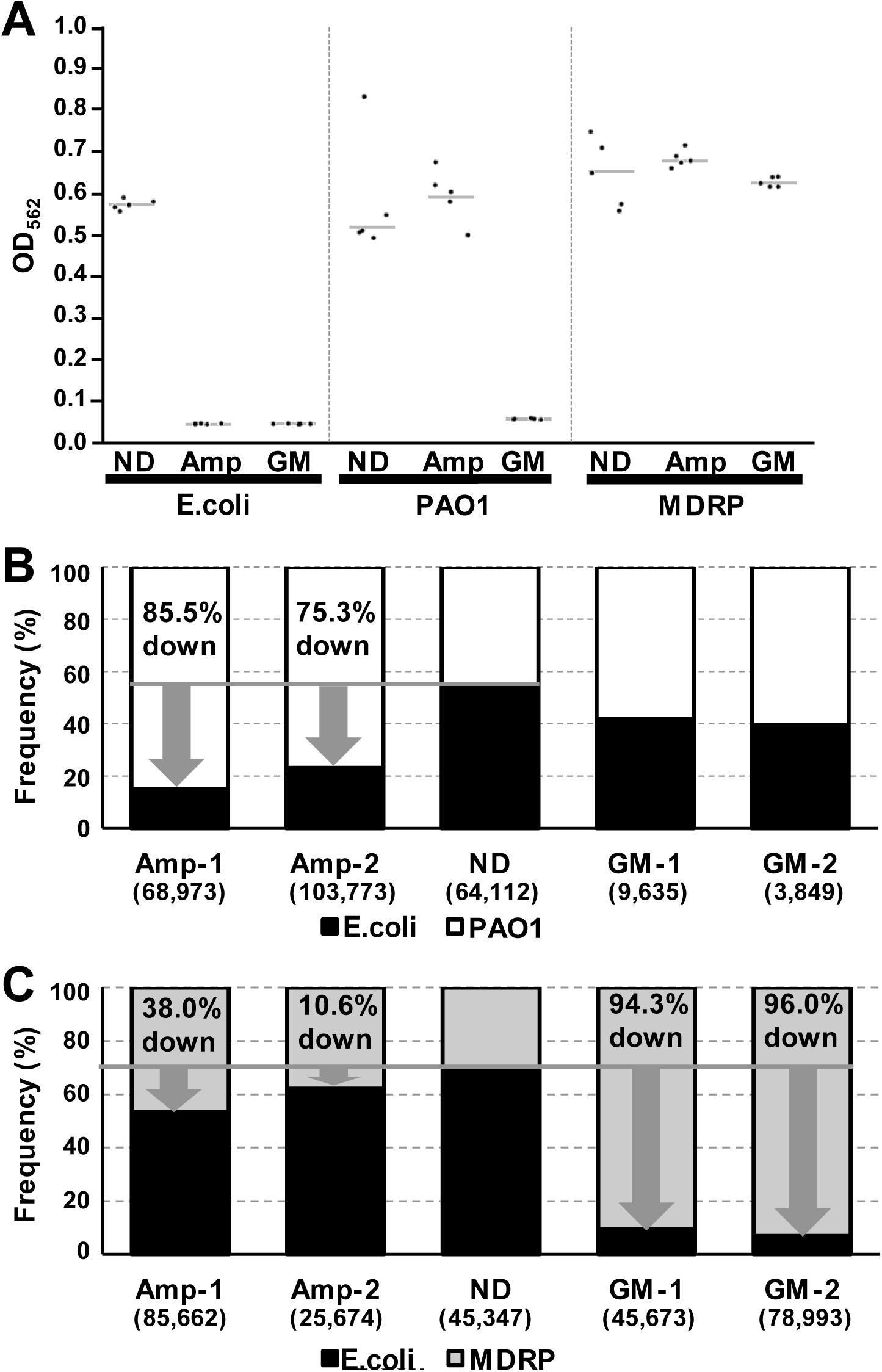
Comparison of bacterial compositions by nanopore DNA sequencing. (A) Results of AST by batch test. The gray lines represent the average OD values. (B) Identification of bacterial compositions by the nanopore DNA sequencer MinION using bacterial mixtures of *E. coli* (black) and PAO1 (white) and (C) *E. coli* (black) and MDRP (gray). The number inside the parentheses is the number of reads. The gray line shows frequency of *E. coli* in untreated bacterial mixture. The arrows show the VBD indeces. The VBD index is the estimated rate of decrease of the number of viable bacteria by the effect of antibiotics. ND, untreated bacterial mixture; Amp, ampicillin treatment; GM, gentamicin treatment.

### Identification of bacterial compositions by 16S rRNA metagenome sequencing

A total of 531,691 reads were obtained in two MinION runs of 16S metagenome analysis. By assigning each read to bacterial species, we calculated the bacterial compositions in each experiment and estimated the effects of antibiotics. In the *E. coli* and PAO1 mixture, the proportion of reads from *E. coli* was drastically decreased when treated by ampicillin (Figure 1B). The frequency of reads for *E. coli* was 54.9% without antibiotic treatment, but it changed to 18% and 22% in two experiments. These correspond to 85.5% and 75.3% decrease of viable bacteria in terms of VBD index. When treated by gentamicin, to which both *E. coli* and PAO1 are sensitive, the proportion of reads from *E. coli* did not decrease drastically. However, from the results of real-time PCR, it was confirmed that the total amount of *E. coli* and PAO1 decreased (Table 2). Next, in the *E. coli* and MDRP mixture, the frequency of reads from *E. coli* dropped to 53.3% and 62.2%, compared to 70.2% without antibiotics treatment (Figure. 1C). This corresponds to 38.0% and 10.6% decrease of VBD index. When gentamicin was used, the frequency of *E. coli* reads dropped to 9.5% and 6.9%, corresponding to 94.3% and 96.0% VBD index, showing drastic decrease.

### Measuring concentrations of viable bacteria by real-time PCR

In order to reinforce the evidence from 16S metagenome sequencing, we conducted real-time PCR on viable bacterial genomes after antibiotics and PMA treatments. Here, we used universal primers to amplify the V2 region of 16S rRNA genes. First, we measured concentrations of bacterial genomes from culture fluids of *E. coli*, PAO1, or MDRP (Table 1). The Ct values of untreated *E. coli* were 20.6Ct and 22.4Ct, but those of ampicillin-treated *E. coli* were 26.4Ct and 27.1Ct, showing that the amount of *E. coli* apparently decreased with ampicillin treatment. The Ct values of gentamicin-treated *E. coli* were 24.4Ct and 22.4Ct, showing a slight decrease from untreated *E. coli*, suggesting a milder effect of gentamicin on *E. coli*. On the other hand, the Ct values showed no difference between untreated and ampicillin-treated PAO1, while the amount of the gentamicin-treated PAO1 apparently decreased. The Ct values showed not much difference between untreated and treated MDRP by ampicillin or gentamicin. These results are mostly consistent with the results of AST (Figure 1A). Also, these results confirmed that PMA is effective in removing dead bacteria and measuring only viable bacteria.

**Table 1.** Comparison of Ct values after antibiotics and propidium monoazide (PMA) treatment. The effect of PMA was confirmed by comparing the Ct values obtained by real-time PCR. The bacterial genome of each sample was prepared to 3 ng and used for real-time PCR. The results show the proportion of viable bacteria in the 3ng.

| **Bacteria** | **Antibiotics** | **Ct** | **Tm (°C)** |
| --- | --- | --- | --- |
| *E. coli* | No Drug | 20.6 | 80.6 |
| *E. coli* | No Drug | 22.4 | 80.2 |
| *E. coli* | Ampicillin | 26.4 | 80.2 |
| *E. coli* | Ampicillin | 27.1 | 80.6 |
| *E. coli* | Gentamicin | 24.4 | 80.2 |
| *E. coli* | Gentamicin | 22.4 | 80.2 |
| PAO1 | No Drug | 17.6 | 80.6 |
| PAO1 | No Drug | 16.4 | 80.6 |
| PAO1 | Ampicillin | 17.2 | 80.6 |
| PAO1 | Ampicillin | 16.0 | 81.0 |
| PAO1 | Gentamicin | 23.8 | 80.6 |
| PAO1 | Gentamicin | 23.0 | 80.6 |
| MDRP | No Drug | 19.2 | 80.6 |
| MDRP | No Drug | 16.5 | 80.2 |
| MDRP | Ampicillin | 19.2 | 80.6 |
| MDRP | Ampicillin | 19.2 | 80.6 |
| MDRP | Gentamicin | 18.2 | 80.2 |
| MDRP | Gentamicin | 18.2 | 80.2 |

**Table 2.** Comparison of Ct values after antibiotic and propidium monoazide (PMA) treatment in bacterial mixtures. The effect of PMA was confirmed by comparing the Ct values obtained by real-time PCR using bacterial mixture. The bacterial genome of each sample was prepared to 3ng and used for real-time PCR. The results show the proportion of viable bacteria in the 3ng.

| **Bacterial mixture** | **Antibiotics** | **Ct** | **Tm (°C)** |
| --- | --- | --- | --- |
| *E. coli* and PAO1 | No Drug | 18.6 | 81.0 |
| *E. coli* and PAO1 | No Drug | 19.6 | 80.2 |
| *E. coli* and PAO1 | Ampicillin | 19.4 | 81.0 |
| *E. coli* and PAO1 | Ampicillin | 18.4 | 81.0 |
| *E. coli* and PAO1 | Gentamicin | 23.4 | 80.2 |
| *E. coli* and PAO1 | Gentamicin | 24.3 | 80.6 |
| *E. coli* and MDRP | No Drug | 17.9 | 81.3 |
| *E. coli* and MDRP | No Drug | 18.2 | 81.0 |
| *E. coli* and MDRP | Ampicillin | 19.1 | 81.0 |
| *E. coli* and MDRP | Ampicillin | 20.3 | 81.0 |
| *E. coli* and MDRP | Gentamicin | 21.4 | 81.0 |
| *E. coli* and MDRP | Gentamicin | 20.5 | 80.6 |

Next, we conducted real-time PCR on genomic DNA extracted from bacterial mixtures after antibiotics and PMA treatments that have been examined by 16S metagenome sequencing (Table 2). In the *E. coli* and PAO1 mixture, the Ct values of the untreated mixture were 18.6Ct and 19.6Ct, those of the ampicillin-treated mixture were 19.4Ct and 18.4Ct, and those of the gentamicin-treated mixture were 23.4Ct and 24.3Ct. Although there was no difference between untreated and ampicillin-treated mixture, the amount of total bacteria clearly decreased by the gentamicin treatment. Additionally, in the *E. coli* and MDRP mixture, the amount of total bacteria slightly decreased under ampicillin and gentamicin treatments, which is consistent with the fact that only *E. coli* is sensitive to these drugs.

## Discussion

In this study, we proposed a new method for rapid identification of drug-resistant bacteria by a combination of 16S metagenome sequencing and the use of PMA and antibiotics, and examined its feasibility. First, we assess the effect of PMA and antibiotics on PCR amplification using three bacterial species, and confirmed that only viable bacteria could be amplified by PCR as expected. Next, by a 16S metagenome sequencing of bacterial mixtures after treatment of antibiotics and PMA, we observed changes of bacterial compositions, from which we estimated the existence of drug-resistant bacteria. Furthermore, we examined the total amount of bacterial genomes by real-time PCR, and confirmed that the amount of drug-sensitive bacteria decreased, which was not apparent solely from the proportion of sequencing reads. From these results, we conclude that we can correctly estimate the drug-resistant bacteria by predicting the bacterial compositions and drug-resistance profiles from the proportion of sequencing reads, with the help of real-time PCRs. It is not difficult to determine a drug-resistant species from a bacterial mixture, if not more than one drug-resistant bacteria exist. Even when there is no drug-resistant bacteria, it is possible to judge from the results of real-time PCR, because total amount of bacterial genomes should be significantly decreased.

However, there may be difficult situations, depending on the bacterial species or kinds of antibiotics. For example, metagenome sequencing results of ampicillin-treated mixture of *E. coli* and PAO1 showed significant decrease of *E. coli* (Figure 1B), but real-time PCR showed that total amount of bacterial genome did not change, although one of the bacteria is sensitive to ampicillin (Table 2). In this case, PAO1 might have grown to some extent in one hour even under the ampicillin treatment. We may encounter such an unexpected result in actual cases, so we need to examine effects of antibiotics on as many combinations of bacterial species and antibiotics as possible before putting this method to practical use.

We previously showed that PCR and library preparation for 16S metagenome sequencing can be done in about 1 hour from a given DNA sample (11), and another 1 hour is required to determine bacterial species computationally for each of 5000 reads using reference genomes of 5850 bacterial species (12). Therefore, considering the time for DNA extraction from bacteria (about 1 hour) and irradiation of blue LED (15 min), detection of drug-resistant bacteria will be possible within 4 hours by our method. Of course, further validation of our method is necessary using other antibiotics and other drug-resistant bacteria. However, our method is scalable, and the automation of the experimental procedure is also possible. We hope that, in the future, our method will be useful as a rapid and efficient way to determine bacterial compositions and AST simultaneously.

## Methods

### Sample/culture preparation

We prepared 3 bacterial species of which antimicrobial spectra are known: *Escherichia coli* (ATCC 25922), *P. aeruginosa* (PAO1), and *P. aeruginosa* MDRP. Those bacteria were cultured in heart infusion broth (HIB) (Thermo Fisher Scientific, Waltham, MA, USA) overnight, and were collected by centrifugation. Then, the bacteria pellet was resuspended with saline. The bacterial suspension was double serially diluted using saline on 96-well plate to measure the concentration of viable bacteria. After dilution, OD_562_ was measured in each well using a microplate reader and the quantity of the bacteria was calculated.

### Batch culture experiments

For AST, a bacterial suspension was prepared at 10^7^ colony-forming units (CFU)/mL by HIB. Ampicillin or gentamicin was added to the wells containing bacteria and cultured at 37°C overnight, and OD_562_ was measured in each well using a VersaMax microplate reader (Molecular Devices LLC).

### Antibiotic treatment, PMA treatment and light-emitting diode irradiation

To examine the drug-sensitivity on the genome-base, each bacterial suspension was adjusted to 10^7^ CFU/mL. Mixtures of bacterial suspensions were also prepared. 100 μL of ampicillin (final concentration: 16 μg/mL) or gentamicin (final concentration: 32 μg/mL) was added to 100 μL of each of bacterial suspensions and bacterial mixtures. After the addition of the antibiotics, the bacterial suspensions were cultured for 1 hour at 37°C and mixed well by vortexing (Figure 2A). A 2.5 mM PMAxx (Biotium, Inc., Hayward, CA, USA) was added to the culture medium, and the blue light-emitting diode (LED) was irradiated for 15 min (Figure 2B). Then, bacterial DNA was extracted from the culture medium using Bactozol^™^ (Molecular Research Center, Inc., Cincinnati, OH, USA).

**Figure 2.**
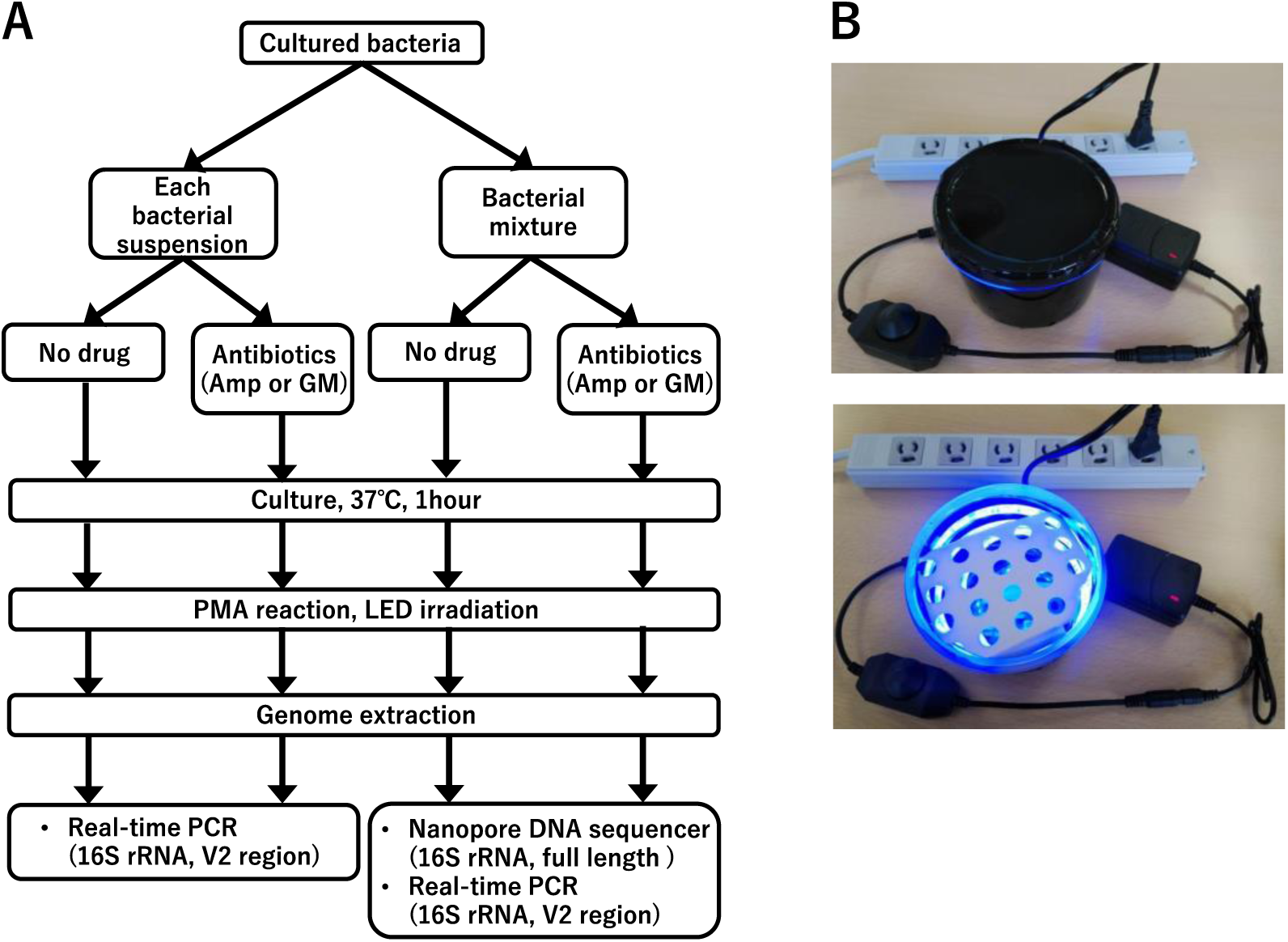
Study design and LED irradiator. (A) Study design. (B) a handmade blue LED irradiator with 60 SMD6060 chips. Top: setup; bottom: when turned on.

### Real-time polymerase chain reaction

To confirm the effect of PMAxx, we used real-time polymerase chain reaction (PCR) targeting the V2 hypervariable regions of the 16S rRNA gene (Figure 2A), using Fast SYBR^™^ Green Master Mix and an ABI 7500 Fast Real-Time PCR System (Applied Biosystems, Foster City, CA, USA).

### Amplification and library preparation

To compare bacteria compositions between samples, a total of 10 ng of bacterial DNA was amplified with the 16S Barcoding Kit (SQK-RAB204; Oxford Nanopore Technologies, Oxford, UK) by PCR as described in the manufacturer’s protocol. Post-PCR clean-up was performed using Agencourt AMPure XP beads (Beckman Coulter, Brea, CA, USA) and eluted in 10 μL of 10 mM Tris-HCl (pH 8.0) with 50 mM NaCl.

### DNA sequencing analysis using MinION

MinION sequencing was performed using the MinION Mk1b sequencer and FLO-MIN106 flow cells. Nucleotides of each read were called by Albacore version 2.1.3 (Oxford Nanopore Technologies), and the sequences were deposited in the DDBJ DRA database (https://www.ddbj.nig.ac.jp/dra/index-e.html) under the accession numbers DRR187692 to DRR187701. Bacterial species were assigned using the minimap2 software **(15)** with the bacterial genomes obtained from the GenomeSync database (http://genomesync.org) as we previously reported **(12, 13)**.

We defined the viable bacteria decrease (VBD) index as the proportion (%) of bacteria killed by the effect of antimicrobial drugs. This can be estimated from the number of sequencing reads. VBD index is calculated by (1 - (*S* x *R*_*0*_) / (*R* x *S*_*0*_)) x 100, where *R* and *S* are the numbers of reads from drug-resistant and drug-sensitive bacteria when antibiotics were used, and *R*_*0*_ and *S*_*0*_ are the numbers of reads from drug-resistant and drug-sensitive bacteria when no drug was used.

## Acknowledgements

We thank the staff of the Department of Laboratory Medicine at the Tokai University School of Medicine for technical assistance.

## Author contributions statement

A.O., K.U., S.A., K.K., S.N., H.M., and T.I. designed the study. S.A. and H.M. collected experimental materials. A.O. and K.U. conducted experiments. A.O., K.K., and S.N. analyzed and interpreted the data. A.O., K.U., S.N., and T.I. wrote the main manuscript text. All authors reviewed the manuscript.

## Additional information Financial support

This work was supported by a grant from the Japan Initiative for Global Research Network on Infectious Diseases (J-GRID) of the Japan Agency for Medical Research and Development (AMED) (JP17fm0108023), a specific grant-in-aid from Takeda Science Foundation, and the Kurata Grants of the Hitachi Global Foundation.

