## Supplementary methods for "Rapid profiling of drug-resistant bacteria using DNA-binding dyes and a nanopore-based DNA sequencer"

#### Sample/culture preparation

*Escherichia coli* (ATCC 25922), *P. aeruginosa* (PAO1), and MDRP were cultured in 30 mL of heart infusion broth (HIB) (Thermo Fisher Scientific, Waltham, MA, USA) overnight. After culturing, bacteria were collected by centrifugation (3,000 ×*g*, 15 min). After the bacteria pellet resuspended with 30 mL of saline, the viable bacteria were collected by centrifugation (3,000 ×*g*, 15 min) and finally resuspended with another 10 mL of saline. Sensitivity Test Broth (Nissui Pharmaceutical Co., Ltd., Tokyo, Japan) was used for the dilution of bacteria. The bacterial suspension was double serially diluted (from $2^{-1}$ to $2^{-9}$) using saline. Then, 50 µL of each diluted suspension was dispensed onto a 96-well plate, and 50 µL of the saline was predispensed onto a 12-well plate and serially diluted. After dilution, $OD_{562}$ was measured in each well using a VersaMax microplate reader (Molecular Devices LLC, San Jose, CA, USA) and the quantity of the bacteria was calculated.

**Batch culture experiments**

A bacterial suspension was prepared at $10^7$ colony-forming units (CFU)/mL by HIB and dispensed at 50 μL onto a five-well plate. Ampicillin (final concentration: 16 μg/mL) or gentamicin (final concentration: 32 μg/mL) was added to the wells containing bacteria and cultured at 37°C overnight, and $OD_{562}$ was measured in each well using a VersaMax microplate reader (Molecular Devices LLC, San Jose, CA, USA).

**Antibiotic treatment**

The bacterial suspension was adjusted to $10^7$ CFU/mL. Ampicillin and gentamicin were prepared at 32 and 64 μg/mL. 100 μL of ampicillin (final concentration: 16 μg/mL) or gentamicin (final concentration: 32 μg/mL) was added to 100 μL of bacterial suspension. Second, the bacterial suspensions were mixed at 50 μL each and added to 100 μL of ampicillin or gentamicin. The antibiotic concentration followed the CLSI guidelines. After the addition of the antibiotic, the bacterial suspensions were cultured for 1 hour at 37°C and mixed well by vortexing (Supplementary Figure 1).

**PMA treatment and light-emitting diode irradiation**

Aliquots (200 μL) of bacterial culture were pipetted into clear microcentrifuge tubes. A 2.5 mM PMAxx working solution was prepared by dilution in sterile water, and an appropriate volume was added to the samples for a final concentration of 25 μM. The concentration of PMAxx (Biotium, Inc., Hayward, CA, USA) may need to be optimized depending on the strain and sample composition. PMAxx was added to the culture medium, and then the tubes were mixed well and incubated for 10 min at room temperature and then subjected to blue light-emitting diode (LED) irradiation for 15 min (465–475 nm) (Supplementary Figure 1).

**Bacterial DNA extraction**

DNA was extracted from the culture medium using Bactozol™ (Molecular Research Center, Inc., Cincinnati, OH, USA). Bactozol enzyme solution (100 μL) was added to the bacterial pellet after the removal of PMAxx including the medium and stored for 30 min at 50℃. Then, 400 μL of DNAzol was added to the lysate and stored for 15 min at room temperature. After the supernatant was removed by centrifugation, 100% ethanol was added to the DNAzol–lysate solution and mixed well by inverting. After centrifugation, the supernatant was carefully removed by pipetting. The DNA pellet was washed with 1 mL of 75% ethanol by vortexing, and the residual ethanol was removed by 20 μL of nuclease-free water.

**Real-time polymerase chain reaction**

To confirm the effect of PMAxx, we used real-time polymerase chain reaction (PCR) and quantified the bacterial genome (Supplementary Figure 1). We performed genome titration of drug-sensitive and drug-resistant bacteria using real-time PCR. The V2 hypervariable regions of the 16S rRNA gene were amplified in a total volume of 20 µL comprising 1x Fast SYBR Green Master Mix containing 1 µM forward primer (5′-AGNGGCGNACGGGTGAGT-3′), 1 µM reverse primer (5′-CGTCCTCCCGTAGGAGTCTG-3′), and 3 ng of bacterial DNA. Real-time PCR was performed using Fast SYBR™ Green Master Mix and ABI 7500 Fast Real-Time PCR System (Applied Biosystems, Foster City, CA, USA) according to the following program: 42°C for 2 min, 95°C for 10 min, followed by 40 cycles of 15 s at 95°C and 60°C for 1 min, followed by two cycles of 15 s at 95°C and 15 s at 60°C as a dissociation step.

**Amplification and library preparation**

Amplification and library preparation were completed following the protocol of SQK-RAB204 (Oxford Nanopore Technologies, Oxford, UK). A total of 10 ng of bacterial DNA was amplified with 16S Barcode in SQK-RAB204 (Oxford Nanopore Technologies) by PCR as described. Post-PCR clean-up was performed using 30 µL of Agencourt AMPure XP beads (Beckman Coulter, Brea, CA, USA) and elution in 10 µL of 10 mM Tris-HCl (pH 8.0) with 50 mM NaCl. After the PCR clean-up, the quality and quantity of the amplicons were measured using a NanoDrop and Qubit fluorometer, respectively (Thermo Fisher Scientific), as per the manufacturer's instructions. All amplicons were mixed with the same DNA quantity as pooled barcoded amplicons, followed by the addition of 1 µL of rapid adapter (SQK-RAB204), and incubated for 5 min at room temperature.

### DNA sequencing analysis using MinION

MinION sequencing was performed using the MinION Mk1b sequencer and FLO-MIN106 flow cells. Nucleotides of each read were called by Albacore version 2.1.3 (Oxford Nanopore Technologies), which were deposited in the DDBJ DRA database (https://www.ddbj.nig.ac.jp/dra/index-e.html) under accession numbers DRR187692 to DRR187701. We identified the bacteria species using the minimap2 software (1) and the reference bacterial genomes obtained from the GenomeSync database (http://genomesync.org) as we previously reported (2, 3).
